# The biological consequences of knockout of genes involved in the synthesis and metabolism of H_2_S in *Drosophila melanogaster*

**DOI:** 10.1101/2024.10.09.617392

**Authors:** V.Y. Shilova, L.N. Chuvakova, D.G. Garbuz, A.P. Rezvykh, S.Y. Funikov, A.I. Davletshin, S. Sorokina, E.A. Nikitina, O. Gorenskaya, M.B. Evgen’ev, O.G. Zatsepina

## Abstract

Here, we describe the effect of knockout (KO) of two key genes (*cbs*, *cse*) responsible for H_2_S synthesis in the transsulfuration pathway as well as the inactivation of sulfurtransferase gene (*dtst1*) encoding single domain rhodanese on *Drosophila melanogaster* genome expression and several life-history traits. The analysis of H_2_S production in the KO flies showed its minimal level in the strain with double KO (*CBS-/-; CSE-/-*) while flies with triple KO (*CBS*-/-; *CSE*-/-; *dTST1*-/-) exhibited higher H_2_S level and improvements in lifespan. The transcriptomic analysis of fly bodies from the double KO strain with maximum disturbances of fitness components revealed a profound increase in the expression of genes involved in the functioning of the excretory system. Besides, double and triple KO flies exhibited drastic changes in Malpighian tubules’ appearance. Inactivation of genes related to H_2_S metabolism resulted in altered expression of numerous loci involved in mitochondrial function. While KO of the *dtst1* gene did not affect the genome expression, the triple KO flies exhibited more changes in the whole-body transcriptome than double KO flies. Reduced fecundity in double knockouts correlates with pronounced changes in ovarian transcriptome data. Surprisingly, the knockout of the *dtst1* gene affected the flies’ memory.

## Introduction

In the last two decades, hydrogen sulfide (H_2_S) the simplest of the thiols found in the cells of different organisms from bacteria to mammals joined the other two well-known gases, carbon monoxide (CO) and nitric oxide (NO), as the third essential gasotransmitter (Abe and Kimura 1996; Liu et al. 2022; Wang 2010). Three enzymes: сystathionine*-*β*-*synthase, cystathionine γ-lyase and 3*-*mercaptopyruvate sulfurtransferase (CBS, CSE and MST) through direct or indirect use of sulfur-containing amino acids are involved in the transsulfuration pathway (TSP) providing the endogenous production of H_2_S in various cells and organs (Filipovic et al. 2018; Kabil and Banerjee 2014; Li and Moore 2008; Rose et al. 2017; Zhang et al. 2020), although a certain proportion of H_2_S is produced via-non-enzymatic reactions (Kolluru et al. 2013; Olson et al. 2013; Yang et al. 2019).

It was demonstrated in numerous studies that H_2_S and other reactive sulfur species or persulfides play a key role in different cellular regulatory processes including inflammation, energy metabolism, neurotransmission, redox signalling etc. (Nishida et al. 2012; Yadav et al. 2016). At the same time using CBS/CSE/3-MST triple KO mice, it was shown that these genes involved in TSP are not major sources of endogenous reactive persulfide production (Zainol Abidin et al. 2023).

In addition to hydrogen sulfide production, CBS and CSE are involved in the metabolism of sulfur-containing amino acids. The KO of these genes is deleterious for an organism because it leads to the accumulation of TSP intermediates, homocysteine and cystathionine, and affects methionine levels as well as the products of the methyltransferase pathway (Carballal and Banerjee 2022)

The significance of TSP genes in the pathophysiology of various diseases and recent advances in the development of efficient therapies targeted toward this pathway have been recently summarized in several brilliant reviews (Kabil et al. 2014; Li et al. 2018; Scammahorn et al. 2021; Zhang et al. 2021).

Importantly, H_2_S also plays a crucial role in various physiological processes related to the functioning of the female reproductive system in mammals, including humans (Pilsova et al. 2024; Sun et al. 2024b). The universality and high conservativeness of the ancient adaptogenic system controlling H_2_S synthesis and metabolism in all organisms from bacteria to humans makes *Drosophila* flies an ideal subject for studying the role of both the whole system and its components in various aspects of life functions, from reproduction and longevity to resistance to bacterial infections and behavior. Previously, using CRISPR-Cas9 technology, we obtained deletions of the *cbs* and *cse* genes and studied the effects of KO of these genes on several vital fitness characteristics of *D. melanogaster* flies (Shaposhnikov et al. 2022; Shilova et al. 2020; Zatsepina et al. 2022). Besides, we described the effects of single and double KO of *cbs* and *cse* genes on the transcriptome of males and females (Zatsepina et al. 2020).

The *3-mst* gene encoding 3-mercaptopyruvate sulfurtransferase represents the third important member of TSP. This gene is also involved in H_2_S production in most eukaryotic organisms including mammals, whereas this gene is absent in Arthropoda (Mathew et al. 2011). 3-MST participates in both detoxification reactions (cyanide transfer) and H_2_S production. This protein containing a tandem rhodanese domain belongs to the superfamily of sulfurtransferases (Cipollone et al. 2007). Another gene belonging to the same family encodes a thiosulfate-sulfurtransferase (TST), known as “rhodanese”. Thiosulfate sulfurtransferase is a mitochondrial enzyme which detoxifies cyanide (CN^−^) by converting it to thiocyanate (Cipollone et al. 2007). Importantly, purified MST and rhodanese (TST) can both detoxify cyanide and produce hydrogen sulfide (Nagahara et al. 1995, Nagahara et al. 2019).

Previously, based on the available sequencing data, we inactivated the *CG12279* gene encoding a protein with a single rhodanese domain in *D. melanogaster* using the CRISPR/Cas9 technology (Zatsepina et al. 2020). Initially, we erroneously identified this gene as *3-mst*. However, subsequent nucleotide sequence analysis demonstrated that the gene we studied (*CG12279*) is orthologous to the human *TSTD1* gene (Melideo et al. 2014; Libiad et al. 2018). There is evidence that TSTD1 in humans is involved in the sulfide oxidative pathway and H_2_S catabolism (Melideo et al. 2014). Besides, this gene probably participates in the modulation of HDL cholesterol and mitochondrial function in mice and humans (Zheng et al. 2021). Based on these data, we designated the *CG12279* gene as *Drosophila tst1 –* “*dtst1*” in this paper.

Since the specific physiological roles for most sulfurtransferases are currently unknown (Libiad et al. 2018), it was interesting to monitor the effect of *dtst1* KO on different physiological aspects of *Drosophila* and to study its possible interactions with the major TSP genes i.e. *cbs* and *cse* producing H_2_S. Thus, in our study, we measured H_2_S levels in flies from the generated KO strains and showed that its synthesis was only slightly impaired in all single KO flies. Herein, we have also demonstrated the pronounced effects of knockouts of the studied genes (*cbs, cse* and *dtst1*) in various combinations on several life-history traits including the lifespan, and reproduction parameters as well as on mitochondria and excretion-related loci expression. Therefore, herein, we took full advantage of the *Drosophila* system to elucidate the biological role of H_2_S.

## Results

### H_2_S production is significantly reduced in the strain with double but not triple knockout

It is currently unclear which physiological processes in *Drosophila* involve sulfurtransferase (*tst*) genes and what their interactions are with key genes of the transsulfuration pathway, i.e. *cbs* and *cse*. In the present study, we combined a strain with an inactivated *dtst1* gene with a strain containing a double deletion of the *cbs, cse* genes by genetic methods and obtained the triple KO flies (*CBS-/-; CSE-/-; dTST1-/-).* Since there are non-enzymatic pathways for H_2_S metabolism (Kolluru et al. 2013), the first step in our work was to investigate whether the synthesis of this transmitter is altered in flies with knockouts of genes involved in H_2_S synthesis and/or metabolism. The efficiency of the generated knockouts has been monitored by the RT PCR technique (Suppl. Figure 1).

It is evident from Fig.1 that in strains with a deletion of the *cse* gene, the level of H_2_S synthesis is comparable to its level in the control flies. In the strains with KO of *cbs* or *dtst1* gene, as well as in the triple KO strain, H_2_S production is slightly reduced but remains at a fairly high level. The lowest level of hydrogen sulfide production is observed in the double KO strain containing the deletions of both *cbs* and *cse* genes. Surprisingly, the H_2_S level in this strain is significantly lower than in the triple KO flies (Fig. 1A). This result suggests that *dtst1* probably participates in the pathway of H_2_S degradation and, hence, when KO of *dtst1* is combined with the deletion of *cbs* and *cse* genes, the level of H_2_S in *Drosophila* body is increased, due to the decrease in the rate of its degradation. Hydrogen sulfide produced in double and triple KO flies is probably synthesized by alternative non-enzymatic pathways (Kolluru et al. 2013).

**Figure 1.**
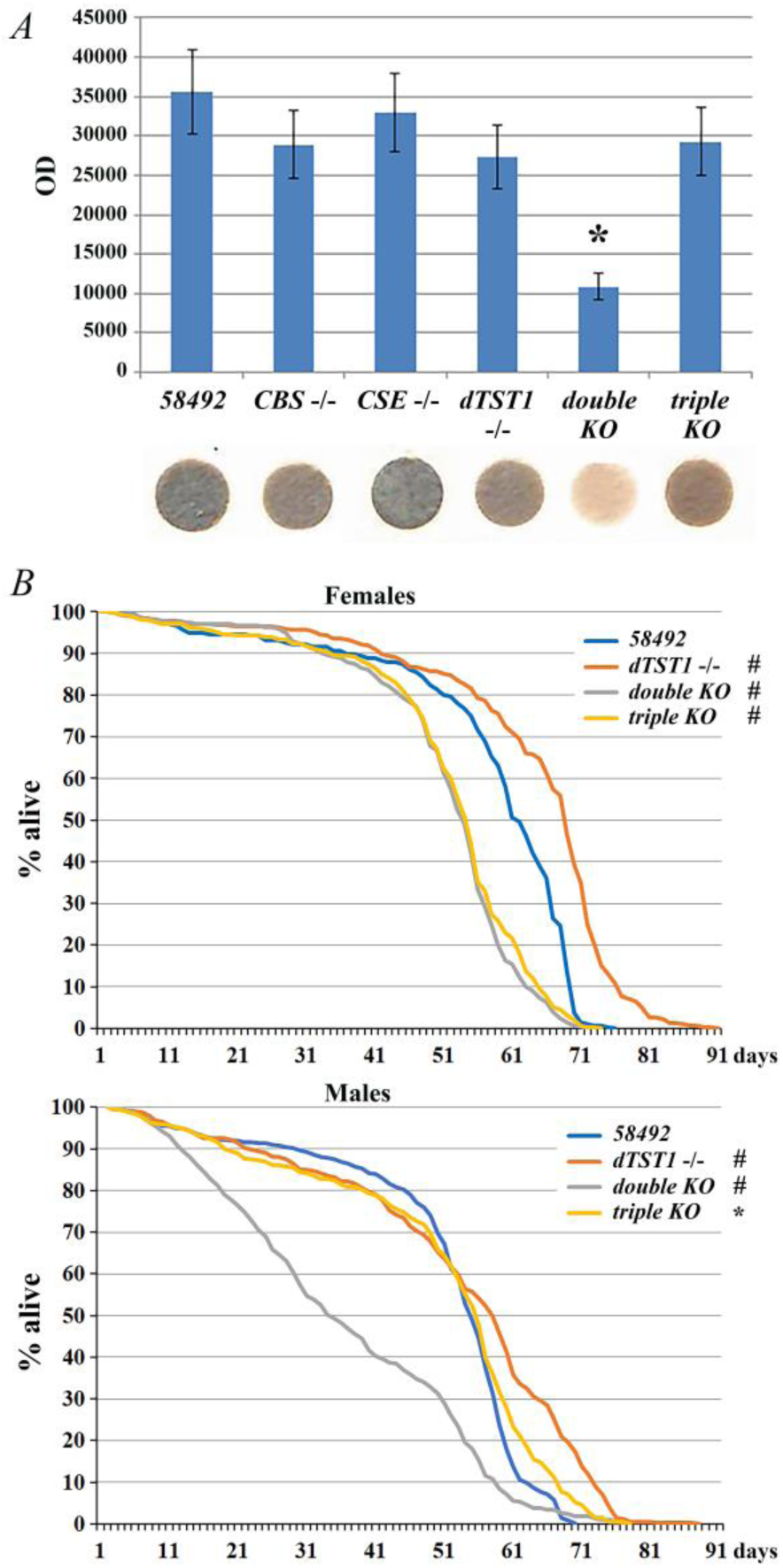
H_2_S production and lifespan of KO strains *(A*) – H_2_S production in extracts from the bodies of seven-day-old *D. melanogaster* flies in the presence of L-cysteine and PLP. H_2_S production was measured using the acetate/lead sulfide method (M@M). Males and females show a similar pattern of hydrogen sulfide synthesis in all strains studied, with females having a slightly higher level of this gas. (*B*) – Effects of *dtst1*KO, and double and triple KO on the lifespan of *Drosophila* males and females * *p* < 0.05, # *p* < 0.001 (v.s. control 58492 flies). Significance was determined by the log-rank test with Bonferroni corrections (lifespan). Lifespan data were pooled from three independent experiments.

The obtained results suggest that in the case of triple KO flies, H_2_S production in the body remains at a rather high level. It is necessary, however, to keep in mind that we measured the levels of H_2_S in the whole body of the flies while different organs and stages of ontogenesis in flies with the studied knockouts may exhibit drastic differences in this parameter.

### Genes involved in H_2_S synthesis and metabolism affect lifespan

Since genes involved in TSP are known to affect multiple major biological systems in animals, it was of interest to investigate how inactivation of individual components of this system, as well as double and triple KO of these genes, affected the lifespan of males and females in the obtained KO strains. It was previously shown (Shaposhnikov et al. 2022) that deletion of the *cbs* gene decreases while deletion of the *cse* gene increases the lifespan of the flies. In the present work, we extended these studies and examined the effects of *dtst1* gene inactivation, as well as the effect of triple KO on the lifespan of *Drosophila*. The original strain *58492*, from which all knockout strains were obtained, was used as a control. The results suggest that KO of *dtst* significantly increases the lifespan of both females and males compared to the control flies. Thus, both the median and maximum lifespan of *dTST1-/-* females is increased by 11% (Supplemental Table. 2) while in the case of males, the effect of *dTST1-/-* is somewhat weaker (Supplemental Table 2). Interestingly, the lifespan of double KO flies, which have the lowest H_2_S levels (Fig.1B), is significantly lower than that of the control flies. Thus, the median and maximum lifespan of the double KO males were 38% and 6.6% lower compared to control males, respectively. In females, this difference is not so pronounced. The median lifespan in double KO females was 9% lower than in the control flies, and the maximum lifespan was 10% lower.

Surprisingly, the introduction of KO of the *dtst1* gene into double KO flies significantly (P < 0.001) shifts the survival curve to the right, increasing the median and maximum lifespan of triple KO males by 64% and 15.8%, respectively. Thus, the survival curve of triple KO males becomes almost identical to that of *58492* males with a slight difference in the late mortality level. However, in the case of females, there are no significant differences in the lifespan between triple and double KO strains, while triple KO females exhibited significantly lower median and maximum lifespan parameters compared to the control flies (Figure 1B). Interestingly, the lifespan of control females as well as females from the *dTST1-/-* and double KO strains is higher than that of the males from the same strain. Thus, the difference in median and maximum lifespan is 13% and 12% for the control strain, 18% and 6.6 for *dTST1-/-*, 38% and 8% for the double KO strain. However, in the case of triple KO strain, the median lifespan of males and females as well as the maximum lifespan are almost the same (Supplemental Table S2).

Major mortality parameters such as mortality doubling time (MRDT), which characterizes the rate of aging, and initial mortality rate (IMR), which is a mortality rate independent of the aging process, were also estimated in our experiments. Characteristically, MRDT values in males of all experimental KO strains are higher than in the control strain (Supplemental Table S2). This indicates that the rate of senescence is significantly reduced in all knockout strains, which may partially explain the observed increased longevity of *dTST1*-/-and triple KO males. However, the increase in mortality in these KO strains takes place earlier than in the control flies due to increased IMR in males of these strains, which mitigates the observed effect of decreased aging rate on the lifespan. Thus, IMR increases most significantly in males from the double KO strain, which affects the shape of the survival curve (Fig.1B). Because of the high contribution of early mortality in double KO males, the curve lost the convex shape characteristic of the control strain and, to a lesser extent, of the *dTST1*-/- and triple KO males.

The differences in MRDT and IMR of females from the KO strains compared to the control strain are not so significant. It is of note, that the IMR of the control females is two times lower than that of males (Supplemental Table S2). In general, the MRDT values estimated in this study do not differ significantly from the corresponding values of most other *D. melanogaster* strains (Promislow and Haselkorn 2002; Shaposhnikov et al. 2018).

### Fecundity changes in the knockout strains

Since fecundity is an important indicator of *Drosophila* females viability we investigated the effects of the generated knockouts on fecundity parameters and monitored the duration of the effective reproductive period in all KO strains in comparison with the control flies.

The analyses performed showed that KO of *dtst1* increased average fecundity by 13% (Fig. 2A, B) while the duration of the reproductive period in this strain was not changed compared to the control strain (Fig. 2A; Supplemental Table S3). Interestingly, double KO flies exhibited a 59% decrease in the number of offspring at the adult stage from a single pair of parents, while triple KO flies demonstrated a 68% decrease in this parameter (Fig. 2A, B; Supplemental Table S4).

**Figure 2.**
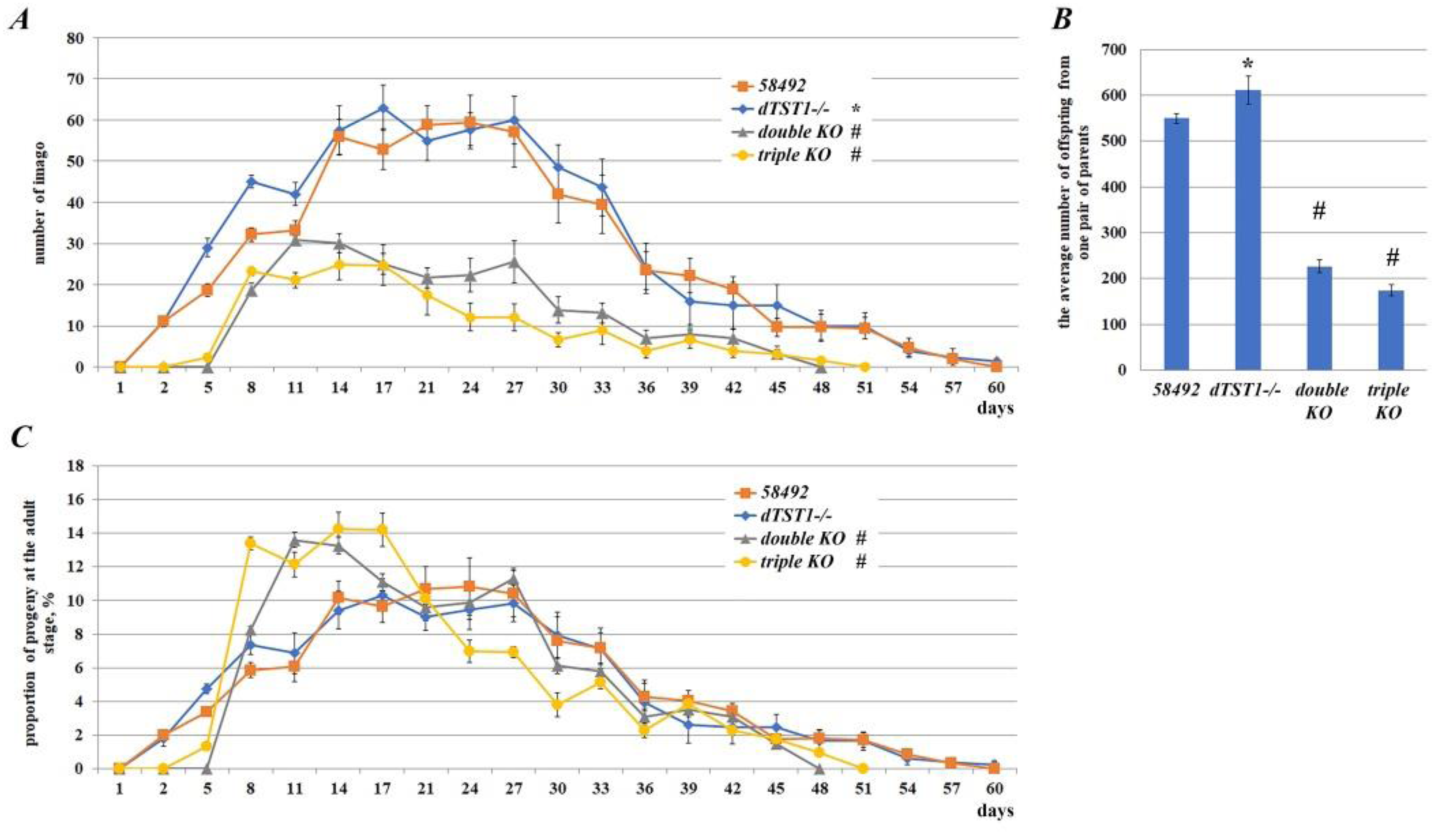
Fecundity analysis of the strains studied. (*A*) – Dynamics of reproductive capacity during the lifespan of one pair of individuals from the strains compared. Eclosed flies were counted every three days (after the second day) *p < 0.05 significance was determined by two-way ANOVA followed by post-hoc Tukey’s HSD tests. (*B*) – The average number of offspring from one pair of parents for the whole reproductive period. (*C*) – The pattern of fecundity rate fluctuations during the lifespan of the compared strains*p < 0.05 significance was determined by two-way ANOVA followed by post-hoc Tukey’s HSD tests

At the same time, the duration of the reproductive period in females from strains with double and triple KO decreased by 16 and 11% (p<0.05) correspondently, compared to the control strain (Fig. 2A; Supplemental Table S3). Figure 2 A, C show that the tested strains differ significantly in reproductive capacity. Thus, flies from the *dTST*-/- strain and the control strain start oviposition somewhat earlier than flies from the double and triple KO strains and significantly exceed them in the number of offspring.

To further characterize the fecundity patterns (Hanson and Ferris 1929) of the analyzed strains, a comparative analysis of the relationship between the age of the parents and the percentage of their offspring (fecundity rate) was performed. The fecundity rate was quantified as the percentage of offspring for every 3 days relative to the total number of offspring (Fig. 2B). The maximum level of fecundity rate was observed in eight- to seventeen-day-old flies from triple KO and eleven-to fourteen-day-old flies from double KO strains. Interestingly, the double and triple KO strains exhibited significantly higher fecundity rates than the control and *dTST1-/-* strains at these time points. In triple and double-KO flies, fecundity begins to decline on the fourteenth – seventeenth day after eclosion, whereas in control and *dTST1-/-* strains it reaches a plateau on the fourteenth day and does not decline until the 27th day after eclosion (Fig. 2B). Thus, the progeny of double and triple KO strains appear predominantly in relatively young flies, which correlates with the observed earlier senescence of these strains (Fig.1B).

To estimate the effect of the obtained knockouts on different developmental stages of *Drosophila*, the number of eggs laid by one pair of flies was counted, followed by the pupae and adult counts (Fig. 3A). Mortality of the flies at embryonic and larval stages was determined as the ratio of the number of pupae to the total number of eggs laid (Fig. 3B). The mortality at the pupal stage was determined as the ratio of the number of the eclosed adults to the number of pupae (Fig. 3B).

**Figure 3.**
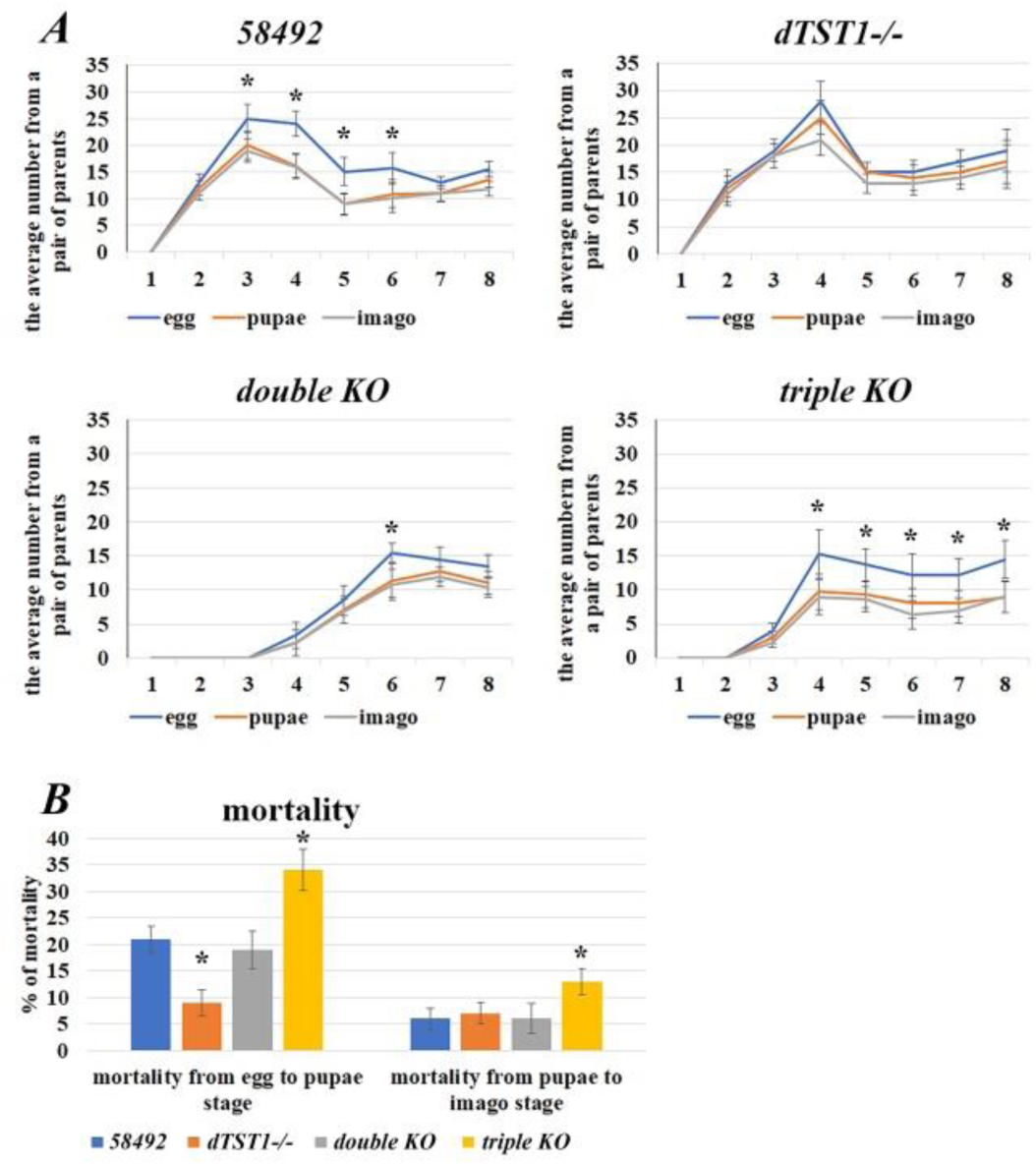
The number of progeny and mortality levels at different stages of ontogenesis of the studied strains. (*A*) – The average number of eggs, pupae and adults from one female when counted daily for eight days. (*B*) – Percent of mortality of tested strains at embryonic, larval (left) and pupal stages (right). p < 0.05, chi-square test with Bonferroni correction.

The lowest mortality at the embryo/larvae stage was observed in the *dTST*-/- strain, the highest in the triple KO strain (Fig. 3B). Similarly, mortality during metamorphosis at the pupae stage was highest (13%) for triple KO flies compared to the control strain (Supplemental Table S5). Thus, surprisingly, the KO of the *dtst1* gene improves several fecundity parameters. However, in the case of triple KO, we observed a significant increase in developmental mortality and reduced reproductive capacity in comparison with double KO and control flies.

### Transcriptome analysis of ovaries of the studied strains

Our experiments revealed the highest level of similar characteristic changes in fecundity parameters in the double and triple KO strains (Fig. 2, 3).

To identify possible causes of the observed interstrain differences, we performed a comparative analysis of transcriptome changes in the whole body and ovaries of double KO strain. Analysis of ovarian transcriptomic libraries allowed us to conclude that as in the case of the whole organisms (Zatsepina et al. 2020), the double KO strain has significantly more genes with altered transcriptional activity compared to strains containing knockouts of single genes involved in H_2_S synthesis and metabolism (i.e. *cbs*, *cse* and *dtst1*) (Fig. 4A). Importantly, the performed analysis of annotated data (https://flybase.org/reports/FBlc0003498.html) demonstrates that *cbs* and *cse* genes are expressed at high levels in *Drosophila* ovaries. Therefore, it is not surprising that the deletion of these genes caused drastic changes in the ovarian transcriptome.

**Figure 4.**
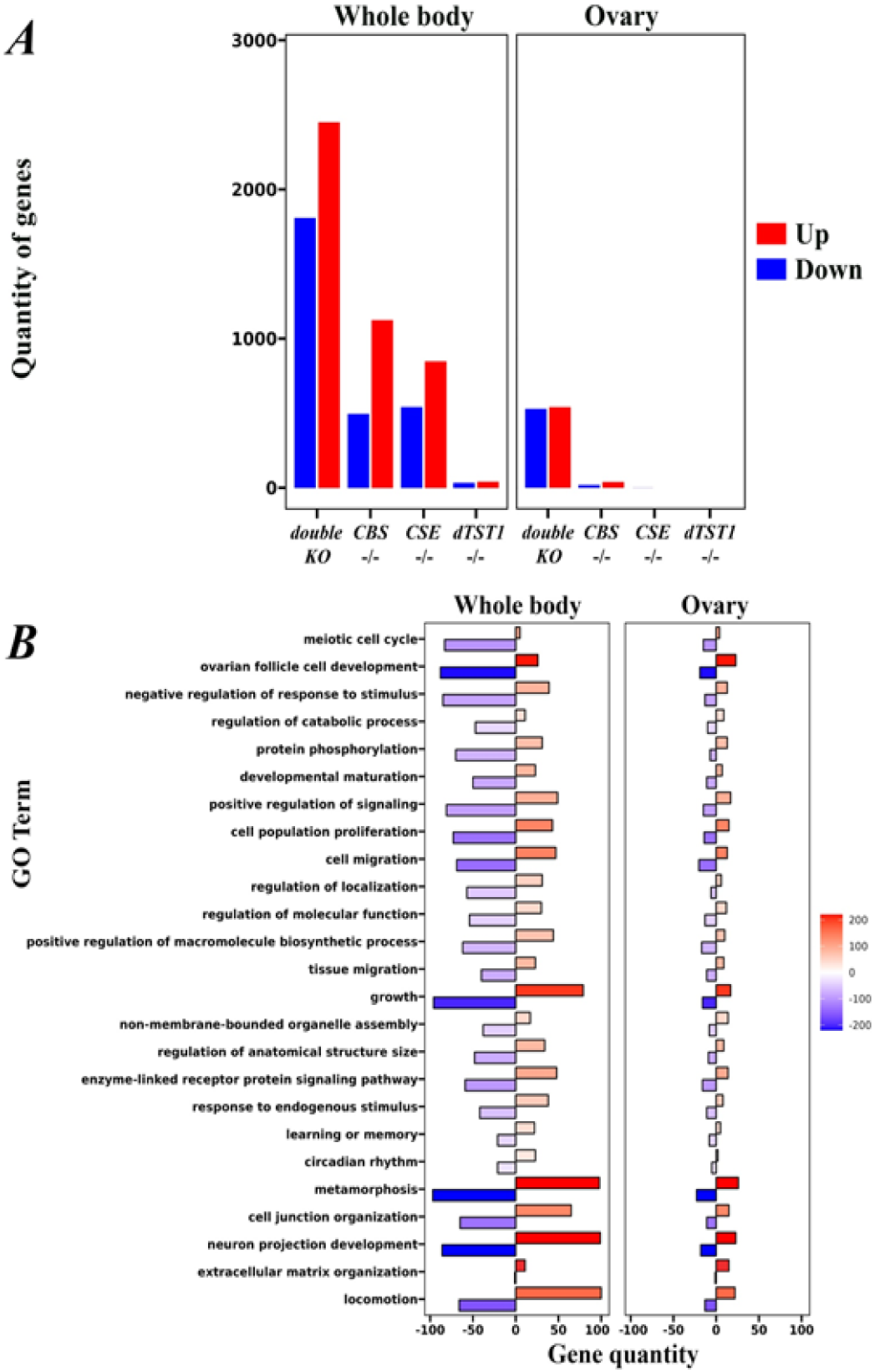
Transcriptome analysis of whole bodies and ovaries of the studied strains. (*A*) – Number of DEGs in the whole bodies and ovaries of flies with single KO of *dtst1, cse, cbs*, and double KO flies compared to 58492 strain (FDR<0.05). (*B*) Gene Ontology analysis of functional groups in the database related to reproduction and development among DEGs in ovarian and whole body tissues in double KO flies. The genes with the decreased expression in ovaries are put to the left, and those increasing their expression level are put to the right. The color scale represents the -Log10 (p-value) of the enrichment test.

The analysis showed generally similar patterns of DEGs in ovaries and whole flies in the strains studied. Moreover, as expected, the number of genes with altered expression in whole flies was several times higher than in the isolated ovaries. However, we revealed an increased proportion of genes with reduced expression levels in the ovaries of double KO flies compared to the whole-body data (Fig. 4A).

Interestingly, the number of genes with altered expression levels in ovaries is higher than in whole flies only for the genes involved in extracellular matrix organization. This group includes genes responsible for the formation of the extracellular matrix, cell structural organization, and basal membrane. The extracellular matrix supports epithelial tissues and is composed of highly conserved components, including collagen IV (Col IV) (Fig. 4B. Supplementary Table S6) (Van De Bor et al. 2021), that are important for *Drosophila* oogenesis and take part in the cell migration, proliferation, elongation, and intercalation processes (Ahmed and Ffrench-Constant 2016; Daley and Yamada 2013; Morrissey and Sherwood 2015). Characteristically, the group of genes with reduced expression levels in whole flies and ovaries comprises genes involved in the meiotic cell cycle. Successful meiosis is necessary to ensure balanced offspring genetics, and the development of viable eggs (Biswas et al. 2021; Hughes et al. 2018; Sun et al. 2024a).

The genes with the most significant changes in expression levels belong to GO categories important for ovarian development and function such as ovarian follicle cell development, growth, metamorphosis and neuron projection development (Figure 4B; Supplemental Table S6 A, B). There is also an imbalance in expression for several other groups of genes in knockout strains compared to control flies (Fig. 4B; Supplemental Table S6).

The demonstrated characteristic changes in the expression of genes involved in ovarian development and function may explain the decrease in the reproductive rates observed in the double and triple KO strains (Fig. 2).

### Changes in the structure and function of Malpighian tubules in double and triple KO strains

It can be seen from the above that deletions of genes involved in H_2_S synthesis and metabolism can affect, to varying degrees, many vital processes and life-history traits in *Drosophila* flies, such as lifespan and fecundity.

Previously (Zatsepina et al. 2020), transcriptomic analysis of the double KO strain showed a significant increase in the expression of several genes controlling the fly excretory system (e.g. *capa, capaR, Uro,* etc.) and genes related to potassium channel function (*Irk3, irk2*, and *KCNQ*). All these genes play an important role in the functioning of the *Drosophila* excretory system, and the observed increase in their expression indicates a disruption of fluid homeostasis and the urgent need for active excretion of toxins and other harmful substances from an organism.

In recent years, the functions of Malpighian tubules (MTs) of *Drosophila* and other insects have attracted much attention from researchers (Dow and Davies 2001; Dow et al. 2015). This organ, which corresponds to the kidneys in mammals (Cohen et al. 2020; Dow 2009; Dow et al. 2022) plays a critical role in the normal metabolism of flies and other insects. Since previously we described the increased expression of several genes involved in excretion processes in KO strains we decided to analyze the structure of the MTs in all these strains. The MTs were stained with the fluorescent dye SF7 to detect differences in the production of hydrogen sulfide. The experiments showed that MTs of all strains actively produce H_2_S (Supplementary Fig. S2).

The performed study of MTs morphology and size allowed us to conclude that in the *dTST1-/-* KO strain, MTs do not differ from those of the control strain. They are thin and long, varying from 25 to 40 µm in thickness and up to 3000 µm in length. In the control strain, the thickness of MTs varies from 30 to 50 μm, and the length varies from 2500 to 3000 μm. Minor vessel thickening is observed in the *CSE-/-* and *CBS-/-* (KO) strains (data not shown) while in the double KO flies the thickness of MTs reaches 45 – 75 μm. The analysis showed that MTs of the triple KO strain exhibited maximum changes in the size and structure (Fig. 5A, Supplementary Fig. S2). Thus the MTs thickness in triple KO flies can reach up to 120 μm, while the total length never exceeds 2000 μm. At the next stage, we investigated the expression of genes involved in the functioning of MTs based on the analysis of transcriptome libraries of five-day females (Fig. 5B). In our comparative analysis of the transcriptome libraries, we explored the available previously described functional transcriptome of *D. melanogaster* MTs (Wang et al. 2004b). It was shown that the KO of the *dtst1* gene does not affect gene expression in MTs (Fig. 5B). At the same time, the combination of this KO with deletions of *cbs* and *cse* genes resulted in triple KO with significantly altered MTs morphology (Fig. 5A) and the expression level of genes related to MTs physiological activity (Fig. 5B).

**Figure 5.**
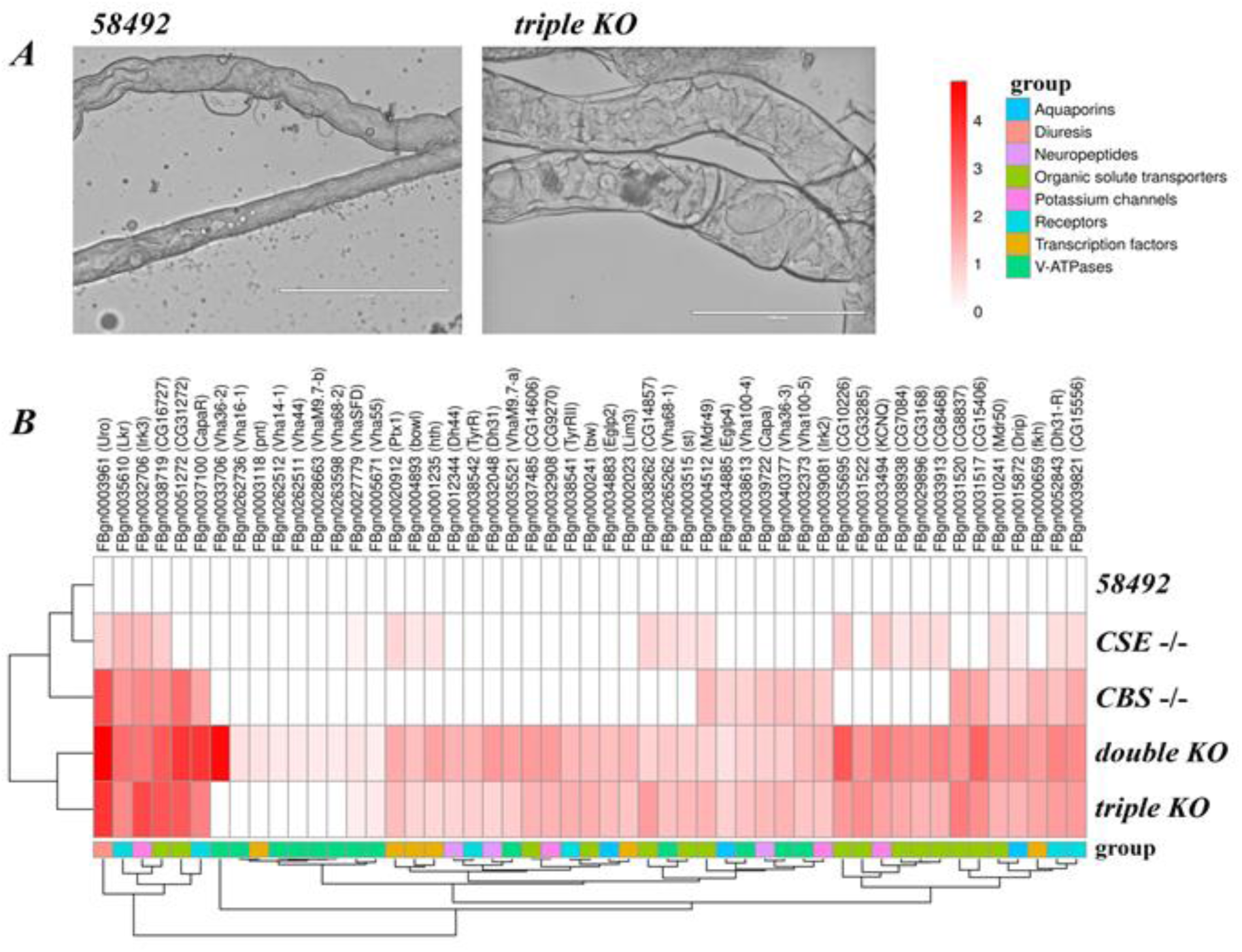
The structure of MTs and expression of pertinent genes in the control and KO strains. (*A*) Photographs of the main segment of MTs from the control and triple KO at the same magnification (10x) were taken using an Evos FL microscope (scale 200μm). (*B*) Heat map of expression of genes related to MTs function in females of the studied KO strains compared to the control strain 58492.

Thus, Fig. 5B shows that in all strains (except *dTST1*-/-) the expression level of *uro* gene, encoding an enzyme localized in MTs and involved in the oxidation of uric acid to allantoin, is significantly increased. A pronounced increase in the expression of several groups of genes related to metabolism and excretion was observed in double and especially triple KO strains. These genes encode receptors localized on the surface of principal cells, i.e. CapaR, DHR31 and peptides binding to them: CAPA, DH31, DH44, acting as diuretic hormones. Diuretic hormones (DH31, DH44) are known to stimulate fluid secretion in MTs by activation of V-ATPases in principal cells of the main segment of MTs (Coast et al. 2001; Cohen et al. 2020). The expression of V-ATPases is predominantly increased in *Drosophila* females with triple and to a lesser extent with double KO (Fig. 5B; Supplemental Fig. S3). Expression of genes encoding receptors on the surface of stellate cells, such as leucokinin and tyramine receptors, is also upregulated in the KO flies. Similarly, the expression levels of several transcription factors that are enriched in MTs, i.e. *pnt, ptx, bowl, hth, fkh, Ets21C, Lim3*, are significantly elevated in double and triple KO flies (Fig. 5B).

The expression level of genes belonging to the superfamily of internal rectifier potassium channels (Kir), which plays an important role in hindgut and MTs osmoregulation (Dates and Kolosov 2024), is also altered in the KO strains.

In all knockout strains, except *dTST*-/-, the expression level of the *drip* gene, which is involved in water transport through the cell membrane, is significantly increased. The expression level of *Eglp4* and *Eglp2* genes involved in the function of water channels is also increased in double and triple KO strains (Fig. 5B).

It is evident from Fig 5B that the expression levels of organic solvent transporters increase differently in strains with deletion of *cbs* or *cse* genes. Thus, genes encoding monocarboxylate transporters such as CG8028 and CG8468 increase the expression level in the strain with *cse* gene deletion, while genes encoding sugar transporters i.e. *CG15406*, *CG31272*, *CG8837*, *CG3285*, *CG14606*, exhibited increased expression in the strain with *cbs* gene deletion. It is of note that the expression of all these genes is significantly increased in triple and double KO strains (Fig. 5B). Supplemental Fig. S3 summarizes the changes in the products of genes with significantly increased expression levels in double and triple KO flies.

### Knockout of dtst1 affects long-term memory

We have previously shown that deletion of the *cbs* gene and double deletion of *cbs* and *cse* genes lead to loss of both short-term and long-term memory in flies, while the deletion of *cse* results in the loss of long-term memory only.

The effect of genes encoding TSTs with a single rhodonase domain on memory has not been previously studied. Since genes with a rhodonase domain, such as *dtst11*, may be involved not only in H_2_S degradation (Melideo et al. 2014) but also in persulfide transport involving partner proteins (Libiad et al. 2018), it was of interest to elucidate the influence of *dtst1*on memory formation processes. Flybase data show that this gene is expressed at a moderate level in the brain of *Drosophila* males (Aradska et al. 2015). Based on these data, we decided to investigate the effect of deletion of this gene on memory in *D. melanogaster*. The study was carried out essentially as before (see M&M for details).

The results obtained allow us to conclude that in males with deletion of the *dtst1* gene, learning indices immediately and 2 days after training remain at a high level and do not differ from the control strain, indicating normal realization of learning processes and formation of short-term memory. However, after eight days, the learning index drops catastrophically (Fig. 6B). Thus, deletion of this gene impairs the formation of long-term memory essentially as described in the case of *CSE-/-* flies (Zatsepina et al. 2022).

**Figure 6.**
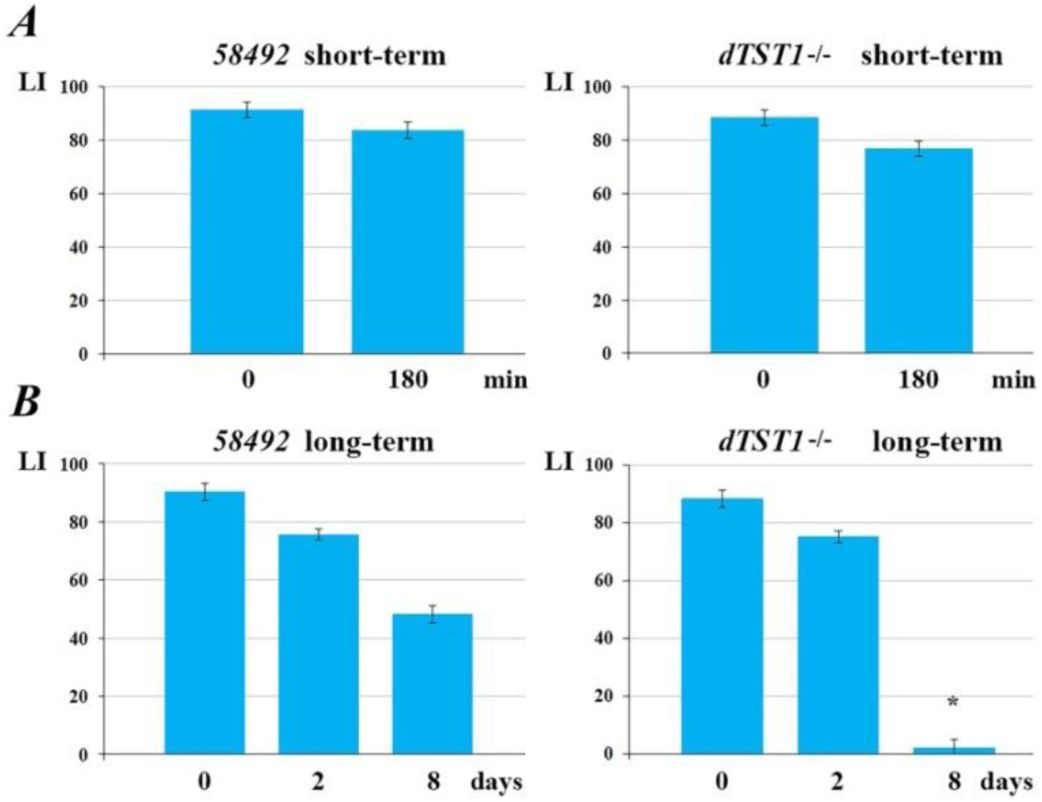
Memory formation in the knockout *dTST1-/-*flies compared with the control strain (58492). (A) – Dynamics of learning acquisition, short-term formation (*B*) term long-term memory acquisition and retention as revealed by conditioned courtship suppression in mutant males. Males of the *58492* and *dTST1-/-* were tested. Abscissa: time after training (min or days); ordinate: LI - learning index, standard units (see M&M). The sample size for each time point was 20 males. * - LI significantly lower than the *58492* strain under similar conditions (two-sided randomization test, α_R_<0.05).

### Changes in mitochondria-related gene expression in knockout flies

Hydrogen sulfide plays a central role in mitochondrial homeostasis and function. It is known that both excess and lack of H_2_S may have deleterious effects at the cellular and organismal levels (Paul et al. 2021). Since double and triple KO flies differ in the level of hydrogen sulfide production it was interesting to find out whether the expression level of genes involved in mitochondrial function is changed in these strains. To reach this goal we analyzed the transcription of genes related to mitochondria function in the studied KO strains (e.g. Krebs cycle enzymes, respiratory chain components, ATP synthase, etc.).

As shown in Fig. 7, a KO of the *dtst1* gene did not result in any changes in DEGs compared to the control strain. Similarly, single deletions of the *cbs* and *cse* genes caused only minor changes in DEGs (14 – 15%), while these changes characteristically differ between these strains. In contrast, double KO leads to dramatic changes in DEGs (i.e. expression level of 57% of genes involved in mitochondria function has been changed). Moreover, the inactivation of all three genes (*cbs, cse,* and *dtst1*) significantly alters the transcription pattern compared to the double KO strain (Fig. 7). It is of note that the observed higher proportion of down-regulated mitochondria-related genes evident in the triple KO flies may represent a compensatory reaction which resulted in the observed higher level of H_2_S in this strain. (Fig.1A). Overall, the expression levels of 70% of the genes involved in the function of mitochondria are altered in the triple KO flies compared to the control strain.

**Figure 7.**
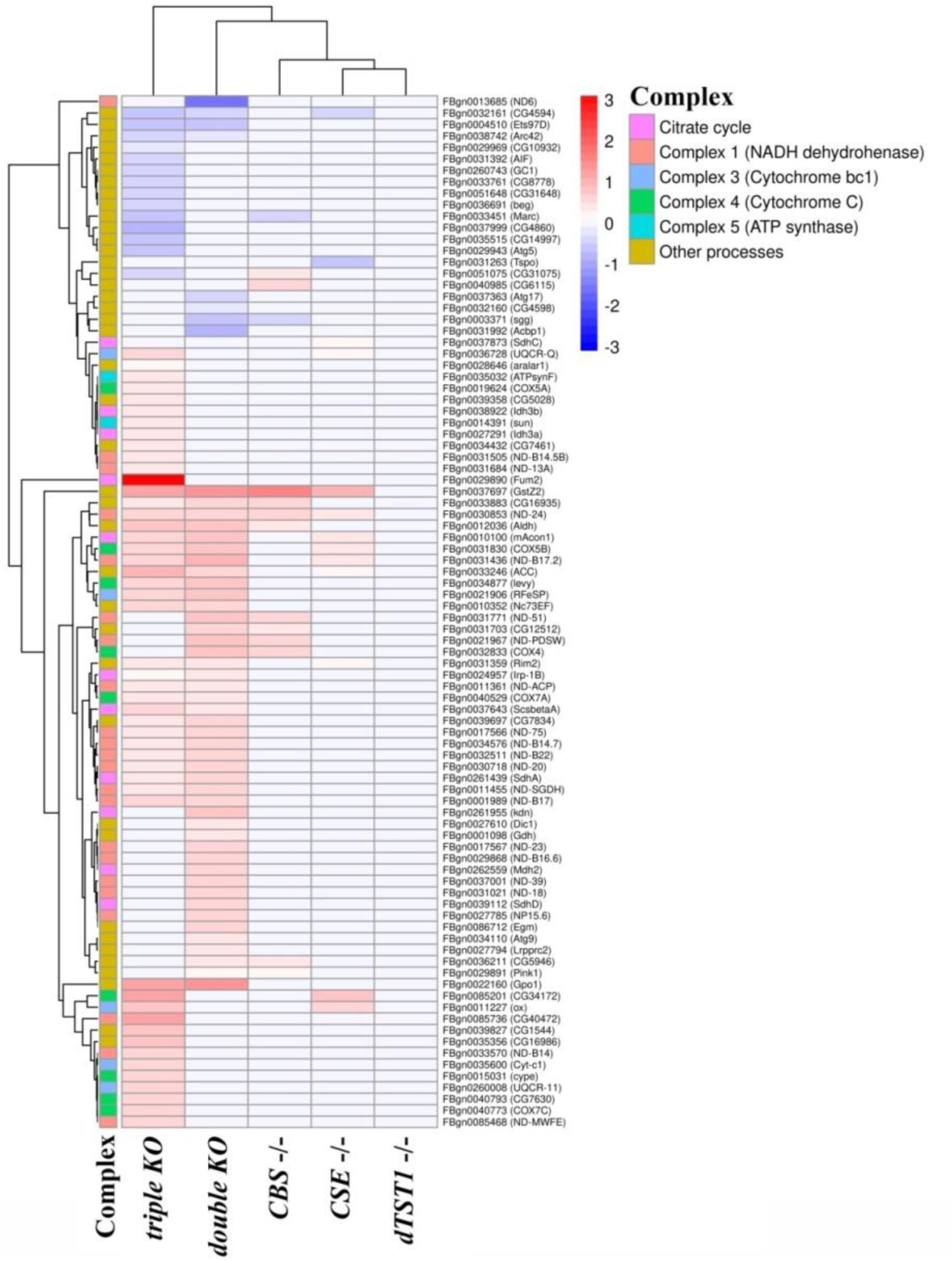
Heat map depicting the expression level of genes involved in mitochondrial function in females from knockout strains compared to control.

## Discussion

The high conservativeness of the ancient adaptogenic system responsible for the synthesis of one of the three major gas transmitters (H_2_S) allows us to consider *Drosophila* flies as a valid and very convenient model for studying the role of H_2_S in virtually all processes associated with life-history traits, including reproduction, aging, immune response, and even mating behavior. Previously, we obtained knockouts of two major TSP genes (*cbs* and *cse),* as well as a third gene (*CG12279*), erroneously named “*3-mst*” in our paper (Zatsepina et al. 2020), using CRISPR/Cas9 technology. We analyzed the effect of deletions of these genes on transcriptomes and some other vital parameters of flies. In the present study, we continued this investigation by obtaining a strain with knockouts of all these three genes, including a third gene (*CG12279*), which, like *3-mst*, belongs to the thiosulfate sulfurtransferase family (TST).

Subsequent bioinformatics analysis of this gene demonstrated that it encodes a protein with a single rhodanese domain. Proteins containing rhodanese domains belong to a wide-spread protein family involved in sulfur transfer reactions between various donor and acceptor molecules and catabolism of H_2_S (Kimura 2020; Kruithof et al. 2020; Mishanina et al. 2015). Importantly, in our case, the studied gene (*CG12279*) encoded a protein with an unusual active site. This site contains a catalytic cysteine followed by a positively charged lysine and a hydrophobic alanine, whereas the catalytic cysteine of canonical TST is followed by two charged amino acids (Libiad et al. 2018) (Supplemental Fig. S4). Notably, the programs MitoFates (Fukasawa et al. 2015), TPpred3 (Savojardo et al. 2015), TargetP 2.0 (Almagro Armenteros et al. 2019) predict that the protein encoded by *CG12279* gene should localize in the cytoplasm and not in the mitochondria which is a characteristic feature of canonical TST family enzymes. TST and MST have similar physicochemical and catalytic properties because they are evolutionarily related enzymes belonging to the same family (Buonvino et al. 2022; Kruithof et al. 2020; Nagahara et al. 1995). The catalytic activity of these two enzymes depends on the cysteine residue in their active site (reviewed in (Kaczor-Kamińska et al. 2021). To this end, it was shown that in the mice knock-out for the *3-mst* gene, the expression of the *tst* gene is compensatory increased (Nagahara et al. 2019).

Since nucleotide sequence analysis shows that the gene we studied (*CG12279*) is structurally homologous to human TSTD1 (Melideo et al. 2014; Nagahara et al. 2019), we designated it as *Drosophila tst1* – “*dtst1*”. It is necessary to emphasize that phylogenetic analysis has shown that arthropods, including *Drosophila*, do not have the canonical *3-mst* gene in their genome (Mathew et al. 2011). It should be mentioned that bioinformatics allowed us to detect three genes with a single rhodanese domain in the *D. melanogaster* genome. Two of them are highly homologous (*CG12279* and *CG4456*) and encode polypeptides of 111 amino acid residues in length. Because the *CG12279* gene, which we named *dtst1*, is expressed at all developmental stages including adult flies, we chose it for further analysis. Interestingly, the sequence of another gene of this family (*CG4456*) in the *Drosophila* genome overlaps with the sequence of the *hsp22* gene and according to the Flybase data, this gene is also stress-inducible and has a similar expression pattern with *hsp22*. The third gene in this family (*CG6000*) differs from genes *CG12279* and *CG4456*. It has a different intron-exon structure and encodes a protein 154 amino acids long. Interestingly, our transcriptomic data show that in triple KO flies, *dtst1* KO causes a compensatory increase in the expression level of the *CG6000* gene, but not of the stress-inducible gene *CG4456* (Supplemental Fig. S5). Thus, we have inactivated a sulfurtransferase gene encoding a protein with a single rhodanese domain that appears to be involved in H_2_S metabolism (breakdown) of H_2_S. This allows us to study the role of sulfurtransferases at the cellular and organismal levels both separately and in complex with the major TSP genes (*cbs* and *cse*) involved in H_2_S synthesis in different organs and at different stages of fly ontogenesis.

It should be emphasized that in our study we observed an increase in H_2_S production in triple KO flies compared to flies with deletion of both H_2_S-producing genes (*cbs* and *cse*) (Fig.1). This suggests that *dtst1* is involved in the process of H_2_S metabolism and degradation since the concentration of this gas in the cells and tissues depends on both the level of its production and the rate of its degradation (Kabil and Banerjee 2014; Vitvitsky et al. 2012).

The knockout of *dtst1* has also an unexpected effect on various fly life-history traits and metabolism. Thus, a single deletion of this gene increases the fecundity of females and causes a significant increase in the lifespan of both males and females (Figures 1, 2). At the same time, our previous transcriptomic analysis (Zatsepina et al. 2020), showed that knockout of the *dtst1* gene *per se* has only a very minor effect on genome expression in the fly body (Fig. 4 A, Supplemental Table S6).

On the other hand, the maximum reduction of lifespan is observed in males of the strain with double KO of TSP genes. Thus, the average lifespan of double KO males was 38% shorter than that of the control strain. Analysis of MRDT and IMR values revealed interesting patterns. Higher MRDT values in males of all knockout strains indicate a decrease in their aging rate. At the same time, double KO males, exhibited significantly elevated IMR values, indicating increased early mortality not related to aging processes but apparently caused by various disorders. This suggests that the double KO of TSP genes which reduces H_2_S production, significantly disrupts homeostasis predominantly in males. Triple KO males show decreased IMR and mortality rates in comparison with double KO males, possibly due to increased levels of H_2_S in this strain leading to partial restoration of homeostasis. However, the lifespan of the triple KO females and their fecundity remain similar to that of the flies from the double KO strain.

It is of note, that we have previously shown that females with double deletion (*cbs* and *cse*) exhibited altered expression levels of significantly more genes than males from the same strain (Zatsepina et al. 2020). The major gene categories that increase the expression level in these females include pathways involved in glutathione metabolism, redox processes, and xenobiotic metabolism (Zatsepina et al. 2020). The increased expression of all these genes likely represents compensation for the knockout of genes responsible for H_2_S synthesis. Importantly, increased expression of many genes from these groups is responsible for longevity (Afschar et al. 2016; Deepashree et al. 2022; Niveditha et al. 2017). Similarly, when gene expression was examined in isolated ovaries, the maximum cumulative effect on transcription of many categories of genes was observed in the double KO strain (Fig. 4).

Characteristically, double KO of the studied TSP genes resulted in a pronounced decrease in fecundity (Fig. 3), while triple KO flies, characterized by comparatively higher H_2_S levels, did not exhibit an improvement in reproductive parameters.

The biological effects of H_2_S are known to be associated with a certain local concentration of this gas in cells and tissues (Li et al. 2011), and although the total H_2_S level in triple KO is higher than in double KO flies, probably its concentration in the ovaries remains insufficient for normal reproduction. It is necessary to mention that the highest level of *cbs* and *cse* expression is observed in *Drosophila* ovaries https://flybase.org/reports/FBlc0003498.html). Similarly, in mice, *cbs* is also strongly expressed in follicular cells at different stages of development and studies of CBS-deficient mice demonstrated that this gene is essential for female reproductive functions in mammals (Guzmán et al. 2006).

In addition, our transcriptomic data demonstrated that double KO flies, exhibit pronounced changes in the expression levels of several groups of genes important for ovarian function and development (Fig. 4, Supplemental Table S6). Moreover, this effect probably is not related to the toxic effect of homocysteine, which should accumulate in the body of flies with *cbs* gene deletion (Škovierová et al. 2016), because these changes are not as significant in *CBS-/-* flies (Fig. 4, Supplemental Table S5).

Our results indicating the participation of the studied genes in the functioning of the excretory system of *Drosophila* are of particular interest. Thus, it was shown that double and triple KOs not only cause a significant increase in the expression of genes responsible for metabolism and excretion but also lead to characteristic changes in the structure and size of MTs (Fig. 5).

It is known that MTs are the most important excretory organs in insects that maintain the correct balance between organic solutes and water (Farina et al. 2022). Interestingly, the KO of the *dtst1* gene alone does not lead to any changes in MTs morphology, but in combination with *cbs* and *cse* deletions, it significantly influences both the appearance of MTs themselves (thickening) and modifies the expression of many genes involved in MTs functioning and detoxification of the whole organism (Fig. 5).

It is known that MTs purify the hemolymph by removing waste metabolites and toxins (Farina et al. 2022). Intermediates of the TSP, i.e. homocysteine and cystathionine, accumulating in the body of double KO flies probably cause intoxication. Thus, intoxication in the strains with *cbs* gene deletion, as well as in double and triple KO flies, is evidenced by an increase in the expression level of the DDT resistance gene *cyp6g1* (Zatsepina et al. 2020), which has previously been shown to be overexpressed in the MTs compared to the entire fly body (Daborn et al. 2002; Joussen et al. 2010). Increased expression levels of ABC-type transporter family genes (mdr50 and mdr49) (Figure 5), which determine multidrug resistance (Seong et al. 2016, Denecke et al. 2022), observed in double and triple knockout strains, are also indicative of an intoxication state of these flies.

Apparently, in the strains with KO of genes responsible for H_2_S synthesis and metabolism, an imbalance of many physiological programs involved in the normal life activity of *Drosophila* takes place. Our comprehensive study showed that in *Drosophila*, as well as in humans and other organisms, genes responsible for the synthesis and metabolism of H_2_S are involved in a wide spectrum of life-history traits. Thus, whereas we previously found that deletions of the major TSP genes (*cbs* and *cse*) impaired male courtship behavior, in this study we demonstrated a significant effect of the *dtst1* gene KO on this parameter. Furthermore, our preliminary data show that all knockouts of genes responsible for H_2_S synthesis and metabolism exhibit a significant effect on many parameters of immunity (paper in preparation).

Numerous studies have investigated the effects of H_2_S on mitochondrial function (Huang et al. 2023; Murphy et al. 2019; Paul et al. 2021). Moreover, a large body of data demonstrated a correlation between mitochondrial misfunction, oxidative stress and aging characteristics across different species (Fridovich 2004; Hekimi and Guarente 2003; Landis and Tower 2005; Wallace 2005).

Similarly, our transcriptomic analysis revealed significant changes in the expression of genes related to mitochondrial function in KO flies. Interestingly, while *dtst1* gene knockout does not affect the expression of these genes when combined with *cbs* and *cse* deletions it causes significant changes in this parameter (Fig. 7). Disturbances in mitochondria functioning may represent one of the main reasons for all sorts of disorders observed in the studied KO strains.

Our studies using strains with KO of the major TSP genes, as well as the gene encoding a single-domain rhodanese involved in H_2_S catabolism, have convincingly demonstrated that this gas transmitter is involved in many physiological processes in the fly. It was also shown that knockout of *dtst1* can both compensate for certain harmful effects of *cbs* and *cse* deletions and significantly enhance their manifestation at the transcriptional and morphological levels.

The generated knockouts of these genes result in severe disruption of homeostasis, which affects all vital processes in the *Drosophila* organism including mitochondria functioning and excretion process. We also hypothesize that deletion of genes involved in the TSP may cause significant amino acid imbalances and changes in fly metabolism that may contribute significantly to all of the abnormalities we observed in the knockout flies. Thus, our study takes a significant step towards understanding the role of genes involved in H_2_S synthesis and metabolism in various life processes, including lifespan as well as reproductive parameters and memory.

## Materials and Methods

### Drosophila melanogaster strains

The development of transgenic knockout (KO) strains bearing CRISPR/Cas9-mediated deletions of cystathionine ß-synthase (*cbs*), cystathionine γ-lyase (*cse*) and *CG12279* genes has been previously described (Zatsepina et al. 2020). Briefly, strain 58492 (BDSC_58492, Bloomington, USA), which was previously used to generate *cbs* and *cse* deletions, was used as a control background strain (Zatsepina et al. 2020). All strains were maintained and experiments were conducted under constant environmental conditions (25°C, 60% relative humidity, 12-h light/dark cycle) that were provided by climate chamber Binder KBF 720 (Binder, Germany).

### H_2_S measurement

Five day-old 20 females or 22 males of each strain were homogenized in a 180 μl PBS containing 10 mM L-cysteine, 1 mM pyridoxal 5’-phosphate hydrate (PLP) and proteases inhibitor cocktail (cOmplete™, EDTA-free Protease Inhibitor Cocktail Roche).

Homogenate was transferred to a 96-well plate. Paper with 2% lead acetate was placed on a top of 96-well plate. Incubation occurs at 30°C for 20 h followed by imaging and analysis of the lead sulfide dots on the paper. The dark circles (dots) are due to the formation of lead sulfide on the filter paper. The optical density of the circles was measured by the ImageJ program.

### Lifespan analysis

Flies from the control and experimental strains were collected in vials on the day of eclosion. Using a carbon dioxide anaesthetizing apparatus (Genesee Scientific, USA) the flies were sorted by sex and placed in vials containing food medium at a density of 25 flies per vial. 12 vials per group were analyzed. The food medium on which the control and experimental flies were maintained during all experiments was prepared according described formulation (Shaposhnikov et al. 2021). The number of dead flies was counted daily starting from the first day of the imago’s life. Food vials were replaced by fresh ones two times per week. All the experiments were performed at least three times. Medium (50%) and maximum lifespan (90% death) were estimated.

Statistical analysis was performed using an online application for survival analysis OASIS 2 (Han et al. 2016). The Kaplan-Meier curve is used to graphically represent the survival function. The log-rank test was used to compare the survival curves. The statistical significance of the differences at the 50th (median lifespan) and 90th percentiles of lifespan was assessed using the Log-Rank test with Bonferroni correction and Wang-Allison (Boschloo’s) test (Wang et al. 2004a), respectively. Coefficients for MRDT and IMR were estimated using STATISTICA software, version 12 (Stat Soft Inc., USA).

### Analysis of the reproductive parameters

For the analysis of fertility dynamics newly eclosed virgin females were individually crossed with two males. All flies of each experimental group consisting of ten females were transferred to vials with fresh medium every 24 h. The number of laid eggs was counted under the SZX10 stereomicroscope (Olympus, Japan). The kinetics of oviposition was estimated as the mean number of embryos produced by a single female per day for the first ten days. After egg counting, vials were kept to determine the proportion of pupated larvae and eclosed flies.

For analysis of the reproductive period, newly eclosed virgin females were individually crossed with two males. Every three days, flies were transferred to a new vial. The number of eclosed flies was counted in each vial during the whole reproductive period.

### The development of triple knockout flies (cbs, cse and CG12279)

To obtain triple KO flies, we used available double KO of *cbs* and *cse* genes and flies from a strain with a knockout of the *CG12279* gene designated in this paper as *“dtst1”* (Zatsepina et al. 2020). At the first stage, we used a balancer strain w, CyO/L; Tb/Sb for a cross with *CBS-/-*; *CSE-/-* to obtain *CBS-/-*; *CSE-/-*; HT/Sb strain. In the case of *dTST1-/-* strain we used balancer strain w; CyO/L; D/Sb and obtained *CG12279-/-*:w; CyO/L strain. At the next stage we combined these strains to obtain triple KO flies with inactivated *cbs*, *cse* and *dtst1* genes. To confirm the inactivation of these genes (*cbs, cse* and *dtst1)* we performed real-time PCR (Supplemental Fig. S1).

### RNA extraction and quantitative real-time PCR

The procedures used were identical to those described in (Shilova et al. 2020). Briefly: total RNA was extracted from adult five-day-old flies or ovaries from control 58492 and knockout strains using guanidine isothiocyanate RNAzol RT (Molecular Research Center, USA) following the manufacturer’s protocol. All experiments were performed with three to five biological replicates and three experimental replicates. The primers used in qRT-PCR experiments are given in Table S1.

### RNA-seq libraries preparation and transcriptomic analysis

Total RNA extraction from whole adult flies and ovaries was performed as described in the previous section. The libraries were prepared and analyzed according to the protocol described earlier in (Zatsepina et al. 2020). RNA sequencing data were uploaded in local FTP server under the URL: http://85.89.102.15:44525/Sulfur_metabolism_RNAseq/. All raw files in FASTQ format as well as tab-separated files containing Flybase gene identifiers and number of read counts spanning in exonic regions of genes are presented. All required information is presented in "metadata.txt" file.

### Test for learning and memory

To evaluate memory formation in *Drosophila* males, we used a conditioned courtship suppression paradigm (CCSP) (Kamyshev et al. 1999). The protocol is fully described in (Zatsepina et al. 2021).

### Malpighian tubules dissection and staining

MTs were dissected from five-day-old females from all investigated strains in PBS and labelled by 2.5 μM SF7-AM in PBS for 30 min for live-cell imaging, washed 3 times with PBS, mounted in PBS and analyzed using an Evos FL microscope at the same magnification of 4x or 10x. MT measurements were performed in the Photoshop program. At least three MTs were analyzed in each strain.

## Author contributions

O.Z., V.S., M.E., designed research, V.S., O.G., S.S., L.C., A.D., S.F., E.N. performed experiments; V.S., L.C., A.R., S.S., O.G., O.Z., D.G. analyzed data; O.Z., S.S., D.G and M.E. wrote the paper.

## Acknowledgements

We are grateful to Drs. E. Zelentsova and D. Karpov for their assistance in the experiments and discussion of the results.

## Funding

This work has been supported by a Grant from the Russian Science Foundation N◦24-14-00216

Appendix A. Supplementary data

